# *Schistosoma mansoni* does not and cannot oxidize fatty acids, but these are used for biosynthetic purposes instead

**DOI:** 10.1101/463984

**Authors:** Michiel L. Bexkens, Mirjam M. Mebius, Martin Houweling, Jos F. Brouwers, Aloysius G.M. Tielens, Jaap J. van Hellemond

**Author notes:** To whom correspondence should be addressed: Erasmus MC, University Medical Center Rotterdam, Dept. Medical Microbiology & Infectious Diseases, PO box 2040, 3000 CA Rotterdam, the Netherlands.

## Abstract

Adult schistosomes, parasitic flatworms that cause the tropical disease schistosomiasis, have always been considered to be homolactic fermenters and in their energy metabolism strictly dependent on carbohydrates. However, more recent studies suggested that fatty acid β-oxidation is essential for egg production by adult female *Schistosoma mansoni*. To address this conundrum, we performed a comprehensive study on the lipid metabolism of *S. mansoni*. Incubations with [^14^C]-labelled fatty acids demonstrated that adults, eggs and miracidia of *S. mansoni* did not oxidize fatty acids, as no ^14^CO_2_ production could be detected. We then re-examined the *S. mansoni* genome using the genes known to be involved in fatty acid oxidation in six eukaryotic model reference species. This showed that the earlier automatically annotated genes for fatty acid oxidation were in fact incorrectly annotated. In a further analysis we could not detect any genes encoding β-oxidation enzymes, which demonstrates that *S. mansoni* cannot use this pathway in any of its lifecycle stages. The same was true for *S. japonicum.* Absence of β-oxidation, however, does not imply that fatty acids from the host are not metabolized by schistosomes. Adult schistosomes can use and modify fatty acids from their host for biosynthetic purposes and incorporate them in phospholipids and neutral lipids. Female worms deposit large amounts of these lipids in the eggs they produce, which explains why interference with the lipid metabolism in females will disturb egg formation, even though fatty acid β-oxidation does not occur in schistosomes. Our analyses of *S. mansoni* further revealed that during the development and maturation of the miracidium inside the egg, changes in lipid composition occur which indicates that fatty acids deposited in the egg by the female worm are used for phospholipid biosynthesis required for membrane formation in the developing miracidium.

## 1. Introduction

The blood dwelling parasite *Schistosoma mansoni* is a causative agent of the neglected tropical disease schistosomiasis that affects over 200 million people worldwide (Colley et al., 2014). Throughout the life-cycle of this helminth, *S. mansoni* encounters various environments and adapts its energy metabolism accordingly. The free-living stages, cercariae and miracidia, live on their endogenous glycogen stores which they completely oxidize to carbon dioxide using Krebs cycle activity and oxidative phosphorylation (Tielens et al., 1991; Van Oordt et al., 1989). Within the mammalian host, adult *S. mansoni* reside in the mesenteric veins, where male and female worms live paired and acquire everything they need directly from the blood of the host. These adult schistosomes have a mainly fermentative metabolism as only a small part of the glucose obtained from the host is fully oxidized to carbon dioxide using Krebs cycle activity and oxidative phosphorylation, while the major part is degraded to lactate (Bueding, 1950; Schiller et al., 1975; van Oordt et al., 1985). Lipids are also obtained from the host. The lipid metabolism of schistosomes is rather compromised, as schistosomes cannot synthesize sterols or free fatty acids *de novo* and must use complex precursors from the host (Brouwers et al., 1997). Until recently it was generally accepted that schistosomes do not catabolize lipids for ATP production. However, in 2009 the genome of *S. mansoni* was published, including an automated annotation which indicated that genes for all enzymes used in β-oxidation of fatty acids are present (Berriman et al., 2009). Automated annotation of genomes is however inherently prone to misannotations and are therefore continuously revised. Currently several of the *S. mansoni* genes earlier annotated as coding for fatty acid β-oxidation enzymes are no longer annotated as such in GeneDB and in databases such as Biocyc.org, WormBase and the KEGG-database.

In 2012 Huang et al. (Huang et al., 2012) reported that oxidation of fatty acids acquired from the host is essential for egg production by female *S. mansoni* worms. Thereafter fatty acid β-oxidation has acquired a lot of attention and most general reports on schistosomiasis now state that fatty acid oxidation is essential for egg production in female schistosomes (Colley et al., 2014; Guigas and Molofsky, 2015; Oliveira et al., 2016; Pearce and Huang, 2015). Subsequently, the postulated fatty acid β-oxidation process was subject of studies on gene expression in schistosomes (Buro et al., 2013; Li et al., 2017) and of studies that aimed to identify novel drugs for schistosomiasis (Edwards et al., 2015; Timson, 2016)

However, the assumption that fatty acid oxidation occurs in schistosomes is still controversial as fatty acid oxidation has never been demonstrated directly in *S. mansoni*, nor in any other parasitic trematode for that matter (Frayha and Smyth, 1983; Rumjanek and Simpson, 1980; Saz, 1981). This has prompted us to perform a comprehensive analysis of the lipid metabolism of *S. mansoni*. We incubated adult worm pairs as well as eggs and miracidia with [^14^C]-labelled glucose and [^14^C]-labelled fatty acids to determine the metabolic fate of these substrates in *S. mansoni*. Furthermore, a genomic analysis was performed to examine the possible presence of genes in the *S. mansoni* genome encoding enzymes involved in oxidation of fatty acids. To further investigate the role of fatty acids in eggs we performed a lipidome analysis of eggs during their development.

## 2. Materials and Methods

### 2.1 Parasites and chemicals

A Puerto Rican strain of *S. mansoni* was maintained in Golden hamsters for which animal ethics was approved (license EUR1860-11709). Animal care and maintenance were in accordance with institutional and governmental guidelines. Adult *S. mansoni* worms were isolated from isoflurane anaesthetized hamsters seven weeks after infection. Worms were collected from the portal vein following heart perfusion with S_20_ medium (20 mM HEPES, 85 mM NaCI, 5.4 mM KCI, 0.7 mM Na_2_HPO_4_, 1 mM MgSO_4_, 1.5 mM CaCI_2_, 25 mM NaHCO_3_ and 20 mM glucose pH 7.4) (Tielens and van den Bergh, 1987)]. *S. mansoni* eggs were obtained from livers of infected hamsters. These livers were homogenized in 1.8% (w/v) NaCl using a MACS homogenizer (Miltenyi Biotec, San Diego, USA). The liver homogenate was treated with 1% trypsin in 1.8% NaCl (BD, New Jersey, USA) for 1 hour at 37° C, where after eggs were isolated by filtration over three sieves with decreasing mesh size (Dresden and Payne, 1981). The eggs were collected and rinsed with sterile water. When indicated, the total isolated egg fraction was separated in immature and mature eggs via Percoll density centrifugation by the method described by Ashton et al. (Ashton et al., 2001). All chemicals used were from Sigma Aldrich, St. Louis, MO, USA unless otherwise specified.

### 2.2 Metabolic incubations

*S. mansoni* worms (10 pairs per incubation) were incubated in 25-ml Erlenmeyer flasks for 2.5 hours at 37°C, 95% O_2_, 5% CO_2_ in 5 ml S_5_^+^ medium (20 mM HEPES, 85 mM NaCI, 5.4 mM KCI, 0.7 mM Na_2_HPO_4_, 1 mM MgSO_4_, 1.5 mM CaCI_2_, 25 mM NaHCO_3_, 5 mM glucose and 1% (v/v) delipidated BSA pH 7.4). All incubations were started with the addition of one of the labelled substrates (all from by PerkinElmer, Boston, MA, USA): D-[6-^14^C] glucose (5 μCi), [1-^14^C] octanoic acid (210 μM, 5 μCi) or [1-^14^C] oleic acid (210 μM, 5 μCi). Radioactive incubations were stopped by acidification of the medium to pH =2 by addition of HCl through the septum of the sealed Erlenmeyer flask. Carbon dioxide was trapped for 1.5 hours in 200 μl 4M KOH in a center well suspended above the incubation medium. Afterwards, the trapping solution was transferred to a vial containing water and Luma-gel (Lumac*LCS, Groningen, The Netherlands), after which radioactivity was measured in a scintillation counter. Worm pairs were removed from the incubation medium and stored at −20 °C until further analysis. The acidified supernatant was neutralized by the addition of 6 M NaOH. The labelled metabolic end-products in the supernatant were analyzed by anion-exchange chromatography on a Dowex 1×8, 100–200 mesh column (Serva) (60 × 1.1 cm) in chloride form (Tielens et al., 1981). The column was eluted successively with 200 ml of 5 mM HCl and 130 ml of 0.2 M NaCl. All fractions were collected and radioactivity was measured in Luma-gel. All values were corrected for blank incubations.

*S. mansoni* eggs (4.3_*_10^4^ – 6.2_*_10^4^) were freshly isolated from infected livers and subsequently transferred to a 25 ml Erlenmeyer flask containing 5 ml of a 1 mM glucose solution supplemented with 100 μg penicillin and 100 units streptomycin per ml. After addition of D-[6-^14^C] glucose (5 μCi) or [1-^14^C] octanoic acid (210 μM, 5 μCi) eggs were incubated for 20 hours at 22°C while shaking gently at 125 rpm. Incubations were stopped and analyzed as described above.

### 2.3 Analysis of incorporated lipids after metabolic incubation

To analyze the incorporation of radioactively labeled fatty acids into complex lipids by *S. mansoni* worms and eggs, lipids were extracted from incubated worms and eggs according to the method of Bligh and Dyer (1959). Prior to the lipid extraction, eggs were disrupted by sonication and adult worms were homogenized by a Teflon-potter. The isolated lipid fraction was subsequently split into neutral lipids and phospholipids by dissolving the total lipid fraction in chloroform after which it was loaded on a 2 ml silica gel 60 column (8 cm tall, 0.5 cm in diameter) equilibrated in chloroform. Neutral lipids were eluted with chloroform followed by elution of phospholipids with methanol. The lipid composition of the neutral lipid and phospholipid fractions was further analyzed by thin layer chromatography (TLC) on Silica G by the methods of Freeman and West (1966) and Skipski *et al*. (1962) respectively. After separation of distinct lipid classes by TLC, the plates were dried and placed in iodine vapor to visualize lipid spots, which were subsequently scraped off. The collected silica was suspended in 1 ml H_2_O and 3 ml Luma-gel was added before radioactivity measured in a scintillation counter.

### 2.4 Lipidome analysis of immature and mature eggs

Approximately 1000 freshly isolated *S. mansoni* eggs were stained with Nile red lipophilic stain (1mg/ml) for 20 minutes, shaking at 1200 rpm at 25 °C in 1 ml 0.9% (w/v) NaCl. After staining, the eggs were washed twice with 1 ml 0.9% (w/v) NaCl and mounted for microscopy. Phase contrast images were captured at 200x magnification. For each bright field image, a corresponding fluorescent exposure was recorded. Eggs were classified by size and developmental stage as described by Jurberg *et al.* (2009).

In order to analyze the lipid content in immature and mature eggs, lipids were extracted as described above. Subsequently, the phospholipid content in mature and immature eggs was quantified by the method of Rouser *et al.* (1970), and the ratio of phospholipids over neutral lipids as well as the lipid species composition was determined by Liquid Chromatography coupled to Mass Spectrometry (LCMS). The extracted lipids were loaded on a hydrophilic interaction liquid chromatography (HILIC) column (2.6 μm HILIC 100 Å, 50 × 4.6 mm, Phenomenex, Torrance, CA) and eluted at a flow rate of 1 mL/min with a gradient from acetonitrile/acetone (9:1, v/v) to acetonitrile/H_2_O (7:3, v/v) with 10 mM ammonium formate. Both elution solutions also comprised 0.1% (v/v) formic acid. The column outlet of the LC was connected to a heated electrospray ionization (HESI) source of an LTQ-XL mass spectrometer (ThermoFisher Scientific, Waltham, MA). Full scan spectra were collected from m/z 450–1050 at a scan speed of 3 scans/s. For analysis the data were converted to mzXML format and analyzed using XCMS version 1.52.0 running under R version 3.4.3 (R-Core-Team, 2016; Smith et al., 2006). Principle Component Analysis provided by the R package pcaMethods (Stacklies et al., 2007) was used to visualize the multidimensional LC-MS data.

### 2.5 Identification strategy to detect genes possibly encoding enzymes required for fatty acid oxidation in the S. mansoni genome

In order to detect genes within the *S. mansoni* genome that are possibly involved in fatty acid oxidation, genes known to be involved in fatty acid oxidation (KEGG pathway 00071) were retrieved from six model reference species: *Caenorhabditis elegans, Crassostrea gigas, Danio rerio, Drosophila melanogaster, Homo sapiens* and *Mus musculus.* The protein sequences of these genes were used as a query in a forward BlastP search against the *S. mansoni* genome with an E-value cut-off of 10^−20^. This forward BlastP search resulted in the identification of 14 *S. mansoni* protein sequences. These proteins possibly involved in lipid metabolism were further investigated by the following annotation strategy. First, using the MUltiple Sequence Comparison by Log-Expectation (MUSCLE) algorithm (Edgar, 2004), the corresponding proteins of these identified *S. mansoni* genes were aligned to the amino acid sequences of the best hit from the forward BlastP query with the six model organisms. Second, these alignments were further investigated to determine whether obvious conserved regions in the proteins of the model organisms are present in the corresponding schistosomal proteins. Third, the *S. mansoni* proteins were used as a query in a reversed BlastP search against the six model organisms to check their identity and to verify if the model organism contains a protein with more similarity to the schistosomal protein than the one used in the forward BlastP where only the proteins involved in fatty acid oxidation were used as query. As a control we also searched in the *S. mansoni* genome for the presence of the enzymes of KEGG pathways 00010 (glycolysis and gluconeogenesis) and 00020 (Citrate Cycle).

## 3. Results

The lipid metabolism of *S. mansoni* adult worms and that of eggs and miracidia was studied by incubations with ^14^C-labelled fatty acids, after which the metabolic fate of these fatty acids was determined by analysis of ^14^C-labelled excreted end products to detect catabolic processes and by analysis of ^14^C-labelled lipids to detect incorporation of fatty acids in anabolic processes.

### 3.1 Lipid metabolism in adult S. mansoni worm pairs

In order to use fatty acids for the production of ATP, fatty acids must be oxidized to carbon dioxide, as fatty acids are too reduced to be fermented. To detect fatty acid oxidation by adult *S. mansoni* worm pairs, the worms were incubated with [1-^14^C] oleic acid, or with [1-^14^C] octanoic acid, a medium chain-length fatty acid that easily crosses membranes. After these incubations we analyzed the formation of ^14^CO_2_. We could not detect any fatty acid oxidation by adult *S. mansoni* worm pairs, as the production of ^14^C-labeled CO_2_ from [1-^14^C] octanoic acid as well as from [1-^14^C] oleic acid was below the detection limit of 0.05 nmol CO_2_ per hour (Table 1). In a simultaneously performed control experiment with [6-^14^C] glucose, approximately 60 nmol CO_2_ was produced per hour by 10 worm pairs, which is comparable to earlier studies (Tielens et al., 1989). This result showed that the parasites were metabolically active and that if production of ^14^C-labelled CO_2_ from fatty acids would have occurred, it would have been detected by our assay system. All together these results showed that the adult worms did not oxidize fatty acids under standard incubation conditions.

**Table 1.** Analysis of end product formation and/or incorporation of labelled substrate in neutral lipids and phospholipids by *S. mansoni* worms or eggs plus miracidia. Analysis of end product formation and/or incorporation of labelled substrate in neutral lipids and phospholipids by *S. mansoni* worms or eggs plus miracidia. Organisms were incubated with labelled glucose or fatty acid for up to 20 hours after which an end product formation was analyzed in the headspace or supernatant of the incubation. *S. mansoni* worms or eggs and miracidia were then analyzed for incorporation of labelled substrate.

|  | End product formation |  |  |  | Incorporation of labelled substrate |  |  |  |
| --- | --- | --- | --- | --- | --- | --- | --- | --- |
|  | CO<sub>2</sub> (nmol/hour) |  | Lactate (nmol/hour) |  | Neutral lipids (nmol/hour) |  | Phospholipids (nmol/hour) |  |
| Substrate | worms~ | eggs +<br>miracidia * | worms~ | eggs +<br>miracidia * | worms~ | eggs +<br>miracidia * | worms~ | eggs +<br>miracidia * |
| [6-<sup>14</sup>C]-Glucose | 62.3 ± 10.4 <sup>a</sup> | 7.13 <sup>b</sup> | 534 ± 74 <sup>a</sup> |  | N.D. | 0.49 <sup>c</sup> | N.D. | 0.19 <sup>d</sup> |
| [1-<sup>14</sup>C]-Octanoic Acid | N.D. | N.D. |  |  | N.D. | 33.81 <sup>e</sup> | N.D. | N.D. |
| [1-<sup>14</sup>C]-Oleic Acid | N.D. | N.D. |  |  | 15.99 ± 3.14 <sup>a</sup> |  | 5.51 ± 2.39 <sup>a</sup> |  |

~ 10 worm pairs per incubation
\* Values are expressed per 50,000 eggs + miracidia
<sup>a</sup> All values represent the mean in nmol per hour per 10 worm pairs and standard deviation of three independent experiments.
<sup>b</sup> Mean of two independent experiments (6.98 and 7.28)
<sup>c</sup> Mean of two independent experiments (0.53 and 0.44)
<sup>d</sup> Mean of two independent experiments (0.19 and 0.18)
<sup>e</sup> Mean of two independent experiments (29.9 and 37.73)
N.D. Not Detectable, below detection limit of assay
empty box not analyzed

Previous research has shown that *S. mansoni* cannot synthesize lipids de novo, and therefore, adult worms take up fatty acids from their environment, after which the fatty acids can be modified by elongation before incorporation in triacylglycerol species and phospholipids (Meyer et al., 1970; Brouwers et al., 1997). To investigate the anabolic fate of the exogenous supplied ^14^C-labeld fatty acids in adult *S. mansoni* worms, we determined the incorporation of [1-^14^C] octanoic acid and [1-^14^C] oleic acid into phospholipids and neutral lipids by adult worm pairs in the above mentioned incubations. This showed that adult worm pairs incorporated oleic acid in both neutral lipids and phospholipids, at rates of approximately 16 nmol and 5.5 nmol per hour per 10 worm pairs, respectively (Table 1).

Incorporation of octanoic acid was not detected (Table 1). From these results and earlier reports (Meyer et al., 1970; Brouwers et al., 1997) it can be concluded that the lipid metabolism of *S. mansoni* adult worms is fairly limited, because adult worms do not *de novo* synthesize nor oxidize fatty acids. Adult schistosomes have to obtain fatty acids from their environment and can modify them, after which they are used as building blocks for the synthesis of phospholipids and neutral lipids, such as triacylglycerol.

### 3.2 Lipid metabolism of S. mansoni eggs and miracidia

We also investigated the lipid metabolism of *S. mansoni* eggs and miracidia by incubation of isolated *S. mansoni* eggs in water with trace amounts of ^14^C-labeled substrates. Although eggs were isolated from liver tissue in 1.8 % (w/v) NaCl and in the dark, a situation in which only a few eggs will hatch and release miracidia (Xu and Dresden, 1990), the subsequent metabolic incubation was performed in water and during the 20 hours incubation a substantial part of the eggs hatched and released miracidia into the medium. Therefore, the results of these incubations reflect in fact the combined metabolic activities of eggs plus miracidia. The *S. mansoni* eggs plus miracidia were incubated with [1-^14^C] oleic acid, [1-^14^C] octanoic acid and with [6-^14^C] glucose. After overnight incubation the production of ^14^CO_2_ was determined, but *S. mansoni* eggs plus miracidia did not produce detectable amounts ^14^CO_2_ from the ^14^C-labelled fatty acids, whereas in the simultaneously performed control incubation with [6-^14^C] glucose 7.1 nmol CO_2_ was produced per hour per 50.000 organisms (Table 1).

Since the supplied ^14^C-labeled fatty acids were not used for fatty acid oxidation by *S. mansoni* eggs and miracidia, we analyzed whether the incubated eggs and miracidia had taken up [1-^14^C] octanoic acid and incorporated it in complex lipids. This revealed that octanoic acid was incorporated in neutral lipids at a rate of 34 nmol per hour per 50.000 *S. mansoni* eggs plus miracidia (Table 1). The in parallel performed incubations with [6-^14^C]-glucose demonstrated that not only fatty acids were incorporated into neutral- as well as into phospholipids, but intermediates of glucose metabolism were also incorporated (Table 1). As *S. mansoni* cannot synthesize fatty acids de novo, the incorporation of ^14^C-label from glucose in lipids, is most likely the result of elongation of fatty acids with acetyl-CoA or of the incorporation of glycerol-3-phosphate as a backbone for triacylglycerol (TAG) or phospholipid biosynthesis. These results showed that *S. mansoni* eggs and/or miracidia take up fatty acids and are capable to incorporate them in complex lipids, but they do not oxidize fatty acids to CO_2_.

To perform a more in-depth analysis of the lipid metabolism of the *S. mansoni* egg during development, freshly isolated eggs were separated in mature and immature eggs (Ashton et al., 2001). Hereafter, we analyzed the distribution and composition of phospholipids and neutral lipids in the immature and the mature egg fraction. Microscopic analysis of Nile red stained *S. mansoni* eggs showed a differential distribution and content of lipids during the development of the egg (Fig. 1). Immature eggs (panels A-D) contained many large lipid droplets, which disappeared during maturation of the egg (Panels G-H), suggesting that the amount of neutral lipids decreased during development. To analyze the changes in lipid composition during maturation in more detail, the lipid composition in the immature and the mature eggs was analyzed by LC-MS. A significant difference in the phospholipid content in immature versus mature eggs was observed, as the phospholipid content almost doubled from 3.2 pmol to 5.2 pmol fatty acids per egg after maturation and the amount of neutral lipids decreased, although this difference was not significant (Fig. 2). Principal Component Analysis (PCA) of the lipidomic data demonstrated that the lipid composition between mature and immature eggs differed substantially (Fig. 3A), as the relative abundance of many different lipid species from all lipid classes differs between mature and immature eggs (Fig. 3B). For instance, the phosphatidylserine species (40:4) and phosphatidylinositol species (38:4) were more abundantly present in immature eggs, whereas the phosphatidylcholine species 34:1 and 36:1 were more abundantly present in immature eggs. However, no significant differences in the overall phospholipid class distribution, fatty acid chain length and degree of unsaturation were observed (data not shown). All together, these results showed that the lipidome of *S. mansoni* eggs changes during egg development.

**Fig. 1.**
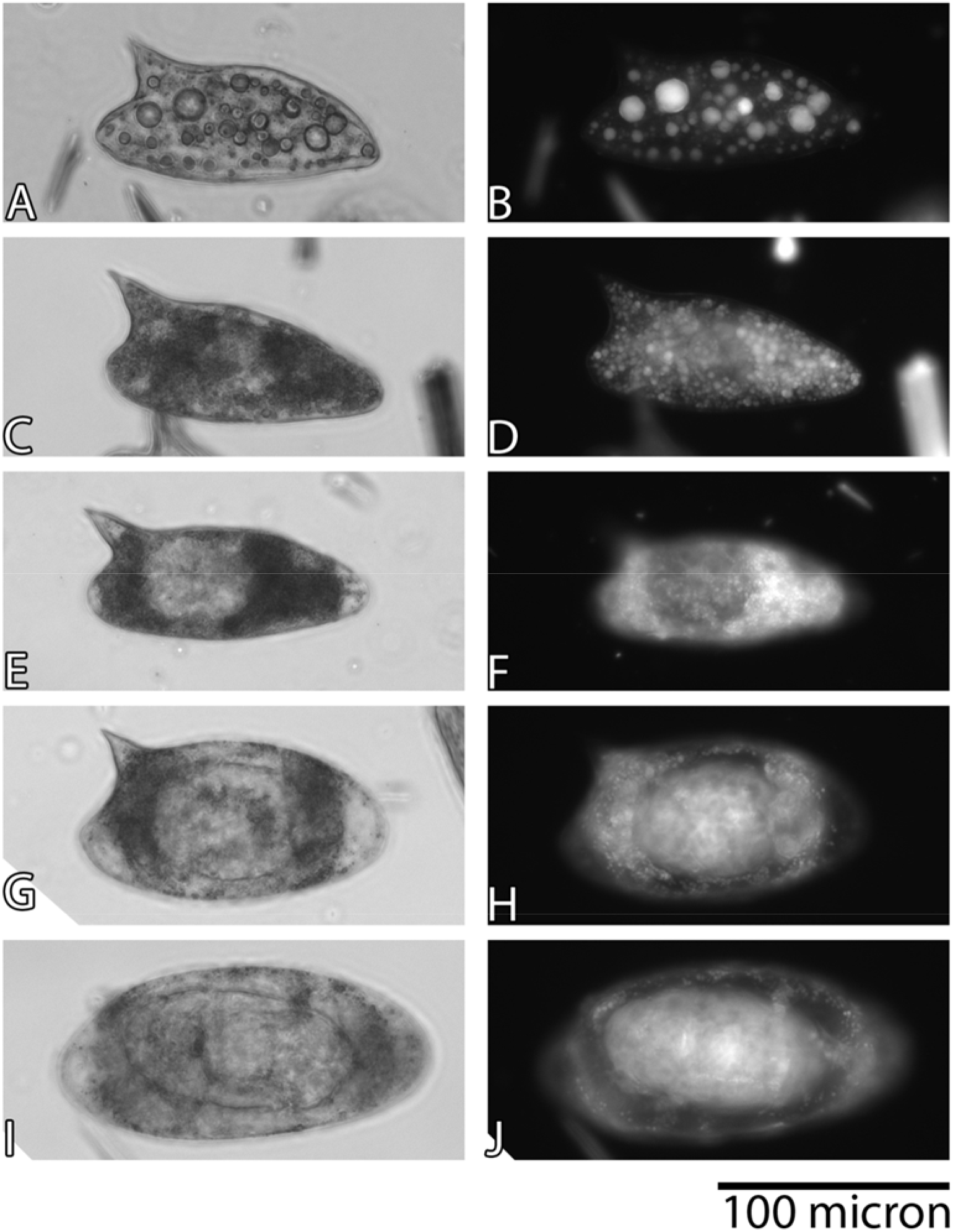
Differential distribution of lipids in developing *Schistosoma mansoni* eggs. Freshly isolated live *S. mansoni* eggs were stained with Nile Red lipophilic stain, after which representative phase contrast images (panels A, C, E, G and I) and their corresponding fluorescent images of Nile Red lipophilic stain (panels B, D, J and H). Eggs are placed in order of maturation; from immature eggs at the top to fully mature eggs with a moving miracidium at the bottom of the figure.

**Fig.2.**
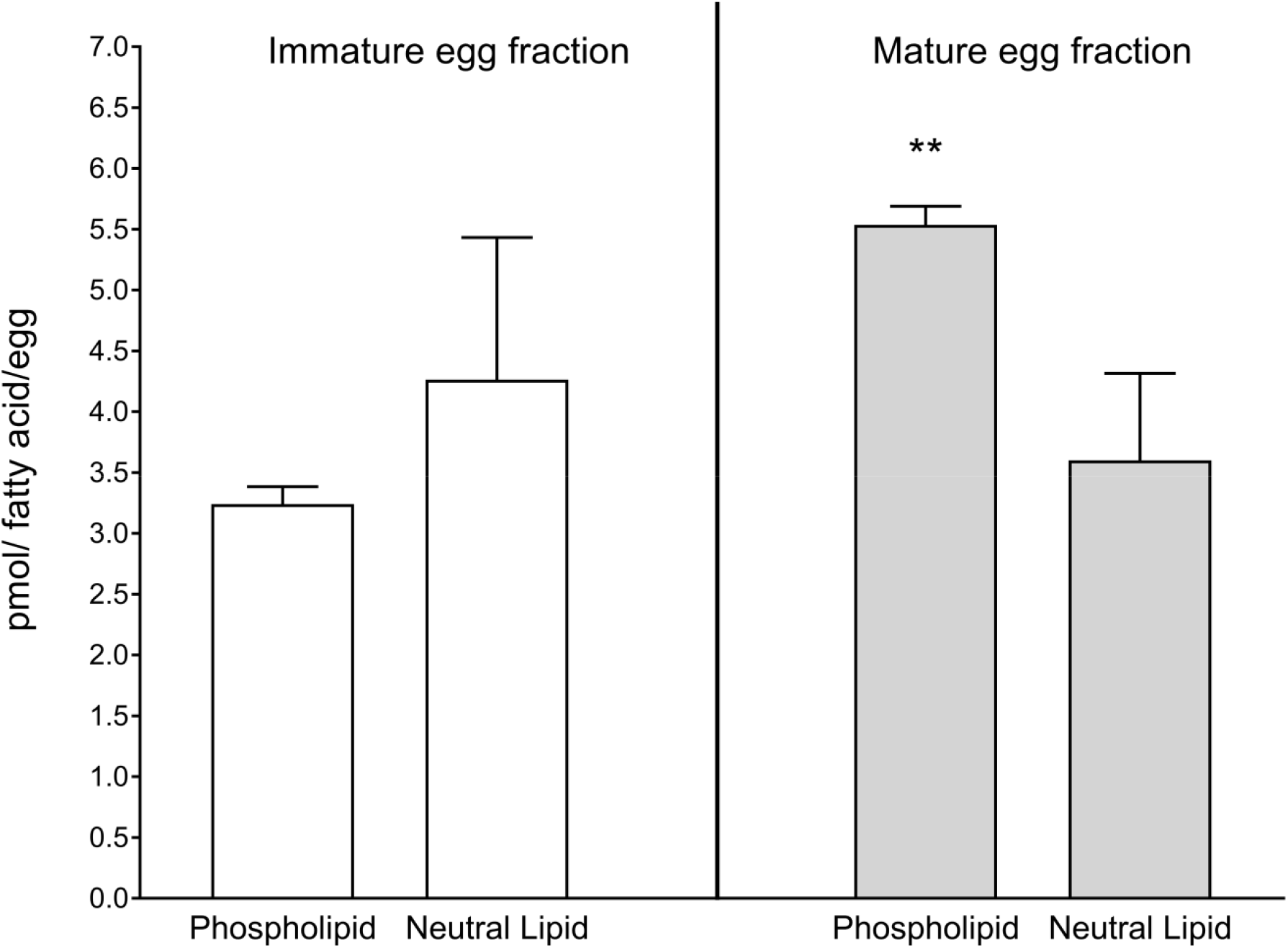
The amount of fatty acids in phospholipids and neutral lipids in *S. mansoni* eggs changes during maturation. The phospholipid and neutral lipid content of *S. mansoni* eggs were analyzed in immature and mature eggs and expressed as picomole fatty acid per lipid fraction. Shown is the average and standard deviation of three independent experiments, each performed in duplicate. ** A one-tailed paired T-test showed a statistically significant increase in the phospholipid content of mature eggs versus immature eggs (P < 0.0054).

**Fig. 3.**
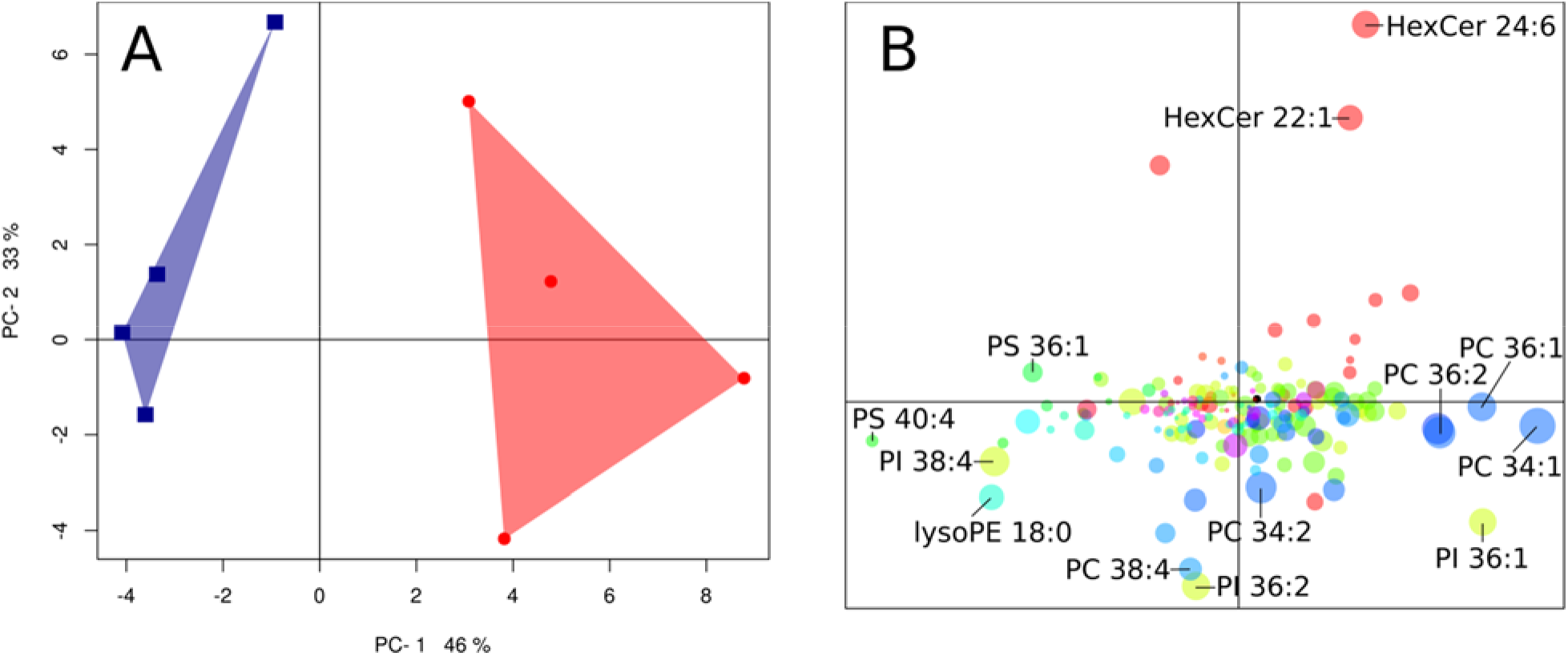
Lipidome analysis of mature and immature *S. mansoni* eggs. Principal Component Analysis on the lipidomic data of mature (blue) and immature eggs (red) revealed different lipid fingerprints for both, as could be concluded from their distinct and non-overlapping score plots (Fig. 3A). The corresponding PCA loadings plot (Fig. 3B) shows that the difference between mature and immature eggs is reflected in many different lipid species from all major phospholipid classes. Abbreviations; HexCer, hexosylceramide; PC, phosphatidylcholine; PE, phosphatidylethanolamine; PI phosphatidylinositol; PS, phosphatidylserine.

### 3.3 Analysis of lipid metabolism at a genomic level

As the investigated stages of *S. mansoni* apparently did not oxidize fatty acids under standard incubation conditions, the question arises whether *S. mansoni* has the genomic capacity to do so at all. To investigate the possible presence of genes in the *S. mansoni* genome which could encode the enzymes required for fatty acid oxidation, we queried the *S. mansoni* genome using genes known to be involved in fatty acid oxidation. Genes annotated in KEGG map00071 "fatty acid degradation" were retrieved from six well characterized model organisms: *Caenorhabditis elegans, Crassostrea gigas, Danio rerio, Drosophila melanogaster, Homo sapiens* and *Mus musculus.* This forward BlastP search resulted in the identification of 14 *S. mansoni* protein sequences. Of the 15 *S. mansoni* proteins, two identifiers were protein isoforms of the same gene, leaving 14 *S. mansoni* genes that might encode enzymes involved in fatty acid oxidation (Table 2). Ten of these 14 genes seemed to encode acyl-CoA synthetases. Of the four remaining genes, three genes seemed to code for homologs of mitochondrial β-oxidation enzymes (acyl-CoA dehydrogenase, enoyl-CoA hydratase and 3-keto-acyl CoA thiolase) and the protein encoded by the last gene seemed a homolog of both carnitine palmitoyl transferase 1 and 2 (CPT-1 and CPT-2). However, the similarity of these last four *S. mansoni* proteins to their corresponding proteins in the six model organisms is probably low, as the E-values were rather high (10^−20^ > E > 10^−80^, Supplementary Table 1). Using this approach, no schistosomal protein was found with a significant homology to enzyme 6 in Fig 4, 3-hydroxyacyl-CoA dehydrogenase.

**Fig. 4.**
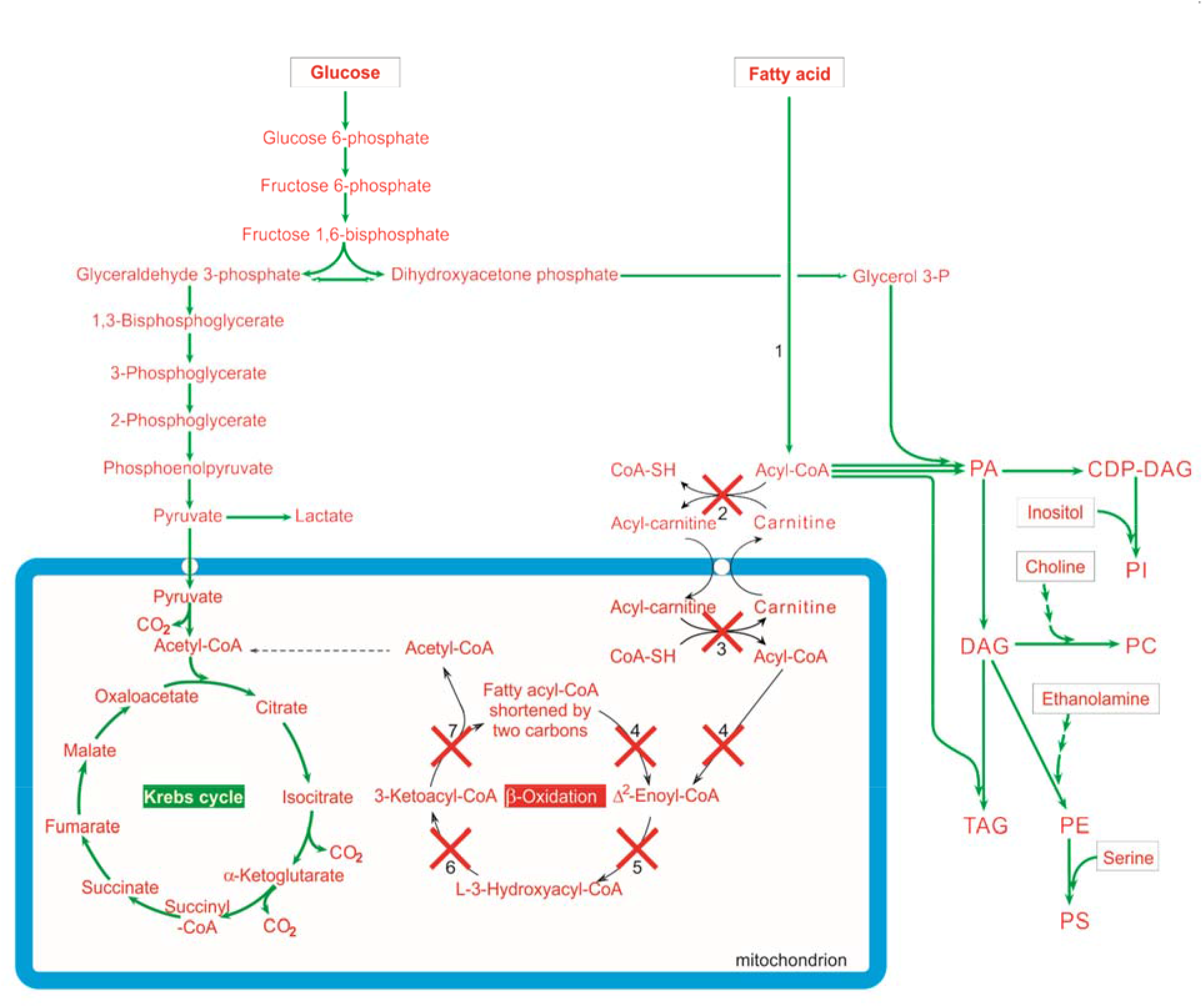
Metabolic map of the glucose and fatty acid metabolism of *Schistosoma mansoni.* Shown are the glycolytic pathway, Krebs cycle, fatty acid β-oxidation and the anabolic processes of triacylglycerol and phospholipid synthesis. Genes encoding the enzymes for glycolysis, Krebs cycle and the anabolic processes are present (green arrows), while genes encoding enzymes for fatty acid β-oxidation are absent in the genome of *S. mansoni* (red crosses). Boxed substrates are supplied by the host. Enzymes: #1. Acyl-CoA synthetase; #2. Carnitine-palmitoyltransferase 1; #3. Carnitine-palmitoyltransferase 2; #4. Acyl-CoA dehydrogenase; #5. Enoyl-CoA hydratase; #6. 3-hydroxyacyl-CoA dehydrogenase; # 7. 3-ketoacyl-CoA thiolase. Abbreviations: CDP]DAG, cytidine diphosphate diacylglycerol; DAG, diacylglycerol; PA, phosphatidic acid; PC, phosphatidylcholine; PE, phosphatidylethanolamine; PS, phosphatidyl serine; TAG, triacylglycerol

**Table 2.** Comparison of BlastP results with genome annotations. Analysis of genes coding for enzymes present in the *S. mansoni* genome. A BlastP analysis was performed using proteins known to be involved in fatty acid oxidation in six model organisms. Results were then ordered by enzyme number 1-7, representing steps in lipid metabolism and beta-oxidation. Identified genes were compared to earlier studies and analyzed using NCBI SMARTblast.

| Enzyme | Enzyme code | Potential homologs identified | Earlier ID (Berriman et al., 2009) | Best hit of reversed BlastP against model organisms | Enzyme code of best hit |
| --- | --- | --- | --- | --- | --- |
| 1. Acyl-CoA synthetase | EC 6.2.1.3 | Smp_103160.1 | Smp_103160 | Acyl-CoA synthetase | EC 6.2.1.3 |
|  |  | Smp_103160.2 |  |  |  |
|  |  | Smp_175090.1 | Smp_175090 | Acyl-CoA synthetase |  |
|  |  | Smp_175090.2 |  |  |  |
|  |  | Smp_175090.3 |  |  |  |
|  |  | Smp_266800.1 | Smp_138660 | Acyl-CoA synthetase |  |
|  |  | Smp_125560.1 | Smp_125560 | Acyl-CoA synthetase |  |
|  |  | Smp_209040.1 | Not mentioned | Acyl-CoA synthetase |  |
|  |  | Smp_165850.1 | Not mentioned | Acyl-CoA synthetase |  |
|  |  | Smp_244300 | Smp_142680, Smp_039870 | Acyl-CoA synthetase |  |
| 2. CPT-1 | EC 2.3.1.21 | Smp_146190 | Not mentioned | choline O-acetyltransferase | EC 2.3.1.6 |
| 3. CPT-2 | EC 2.3.1.21 |  |  |  |  |
| 4. Acyl-CoA dehydrogenase | EC 1.3.8.- | Smp_122930 | Smp_122930 | acyl-CoA dehydrogenase family member 9 | EC 1.3.99.- |
| 5. Enoyl-CoA hydratase | EC 4.2.1.17 | Smp_024930 | Smp_024930 | enoyl-CoA hydratase domain-containing protein 3 | - |
| 6. 3-hydroxyacyl-CoA dehydrogenase | EC 1.1.1.35 | * | Smp_151030 | - | - |
| 7. 3-ketoacyl-CoA thiolase | EC 2.3.1.16 | Smp_267310 | Smp_197330 | acetyl-CoA acetyltransferase, cytosolic | EC 2.3.1.9 |

\* no gene found with significant homology ( $E > 10^{-20}$ )

To further investigate whether or not the retrieved schistosomal genes really encode the corresponding enzymes involved in fatty acid oxidation, we aligned the found schistosomal proteins with the best hit of each model organism (Supplementary Fig. 1.1-1.7). The alignments for the identified *S. mansoni* genes possibly encoding acyl-CoA synthetases (Fig. 4, enzyme number 1) indicated that these *S. mansoni* proteins are true homologues of acyl-CoA synthetases, because the identified *S. mansoni* proteins were highly similar to the ones that encode acyl-CoA synthetases in the six model organisms (Supplementary Fig. 1.1). However, the alignments of the other five retrieved proteins showed low homology to the corresponding proteins of the six model organisms and regions highly conserved in the proteins of the model organisms seemed to be missing in the schistosomal proteins (Supplementary Fig. 1.2-1.7).

We further analyzed these alignments to investigate whether known conserved regions in the proteins in question are indeed absent in the corresponding schistosomal proteins. To this end we produced aligned barcodes of each enzyme where a black bar is produced only when an amino acid is identical in *all* six model organisms and these bar codes were then compared to the found schistosomal proteins (Fig. 5). This analysis clearly showed that the schistosomal acyl-CoA synthetases are indeed highly similar to the ones of the model organisms, as their barcodes are real look-a-likes of the ones of the model organisms. In contrast, the 100% conserved regions in the model organisms of the other five proteins are not mirrored in the schistosomal ones, despite the large gaps the algorithm introduced in three schistosomal proteins (2-4) to produce the best alignment.

**Fig. 5.**
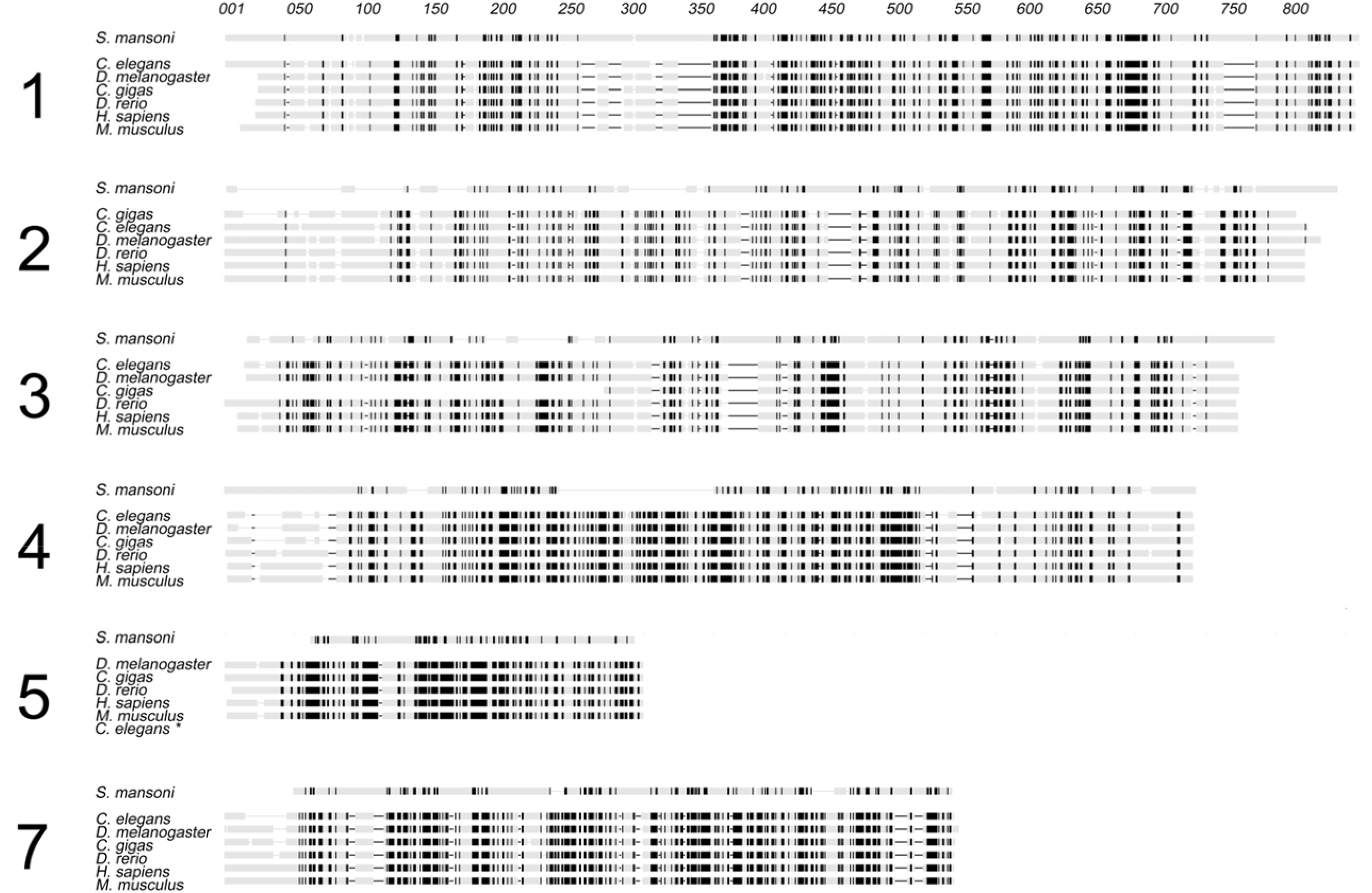
Barcode alignment of *S. mansoni* proteins against proteins involved in fatty acid oxidation. Conserved residues (100% identity) present throughout all six model organisms have been marked as black bars after which it was analyzed whether these residues were also identical in the most similar *S. mansoni* sequences. The resulting barcode shows conserved domains present or absent in *S. mansoni* proteins. The large numbers on the left correspond with the numbering used in figure 4 and table 2. For enzyme number 6, 3-hydroxyacyl-CoA dehydrogenase, no protein with significant homology was detected within the *S. mansoni* genome. The complete alignment is shown in supplemental figure 1. * no gene found with significant homology.

As a final check in this genomic analysis, we performed a reverse BlastP query where the found schistosomal proteins were blasted back against the six model organisms to investigate if the model organisms contain a protein with a higher similarity to the schistosomal protein than the protein used in the forward BlastP where only the enzymes involved in fatty acid oxidation were used as query. This reverse BlastP search revealed that the presumed schistosomal acyl-CoA synthetases show indeed the best homology to the acyl-CoA synthetases of the model organisms (Table 2). However, this reversed BlastP analysis also revealed that the presumed homologs of three β-oxidation enzymes and CPT-1 and CPT-2, were in fact coding for other proteins, or were at least more similar to those other proteins than to the enzymes involved in fatty acid oxidation (Table 2).

As a positive control, the same genomic analysis was performed for KEGG pathway 00010 (glycolysis and gluconeogenesis) and KEGG pathway 00020 (Citrate Cycle), pathways that are both present in *S. mansoni*. This analysis demonstrated that genes could be detected in the *S. mansoni* genome for all enzymes of both pathways. In addition, the identified *S. mansoni* genes were highly homologous to their corresponding genes in the six model organisms, as demonstrated by the low E-values (E<10^−80^, not shown).

All together the results of this genomic analysis showed that the *S. mansoni* genome contains genes coding for acyl-CoA synthetases, which are enzymes that activate fatty acids by coupling them to coenzyme-A (Fig. 4, enzyme number 1). The resulting acyl-CoAs are the beginning of several anabolic pathways that are known to be present in *S. mansoni* (Fig.4). The genome of *S. mansoni*, however, does not contain genes with a decent homology to genes encoding enzymes for fatty acid β-oxidation in the six model organisms. In addition, no genes were found for the two enzymes required for import of fatty acids into the mitochondrion: the cytosolic CPT-1 and mitochondrial CPT-2. We performed this analysis also on the published whole genomes of *S. japonicum, S. haematobium, S. mattheei, S. rodhaini, S. curassoni* and *S. margrebowiei*, which this led to the same result: also all those parasites do not possess enzymes for fatty acid β-oxidation (not shown).

## 4. Discussion

In this study we performed a comprehensive analysis of the lipid metabolism of *S. mansoni* with a special focus on the debated presence or absence of fatty acid β-oxidation in adult worms^1^. Metabolic incubations with the [^14^C]-labelled fatty acids octanoic acid (C8:0) and oleic acid (C18:1) were used to examine the metabolic fate of fatty acids taken up by adult worms and eggs plus miracidia. In addition to a physiologically relevant fatty acid (oleic acid), octanoic acid was studied as well, because of its high bio-availability as it is a medium-chain fatty acid that can pass cellular membranes relatively easy and independent of carnitine shuttles (Schönfeld and Wojtczak, 2016). These experiments showed that fatty acids are taken up from the environment and are subsequently incorporated in complex lipids, such as triacylglycerol and phospholipid species, which is an observation that is in agreement with multiple earlier publications (Meyer et al., 1970; Saz, 1970; Rumjanek and Simpson, 1980; Frayha and Smyth, 1983; Brouwers et al., 1997). Despite the high sensitivity of our assay in which radioactive fatty acids were used, we could not detect the production of carbon dioxide from fatty acids neither by adult worms nor by eggs and miracidia. This showed that these *S. mansoni* stages do not oxidize fatty acids, which confirms an earlier study which demonstrated that miracidia consume their glycogen reserves but not their endogenous triacylglycerol stores (Tielens et al., 1991).

As our present study showed that fatty acids were not oxidized, we re-examined whether schistosomes possess the necessary enzymes to do so. In the first automated genome annotation in 2009 it was reported that the genes for the enzymes required for fatty acid β-oxidation are present in the genome of *S. mansoni* (Berriman et al., 2009). Our analysis to resolve the possible presence of genes coding for fatty acid β-oxidation enzymes in *S. mansoni* demonstrated that the genes coding for enzymes required for fatty acid β-oxidation are absent in the *S. mansoni* genome. This discrepancy with earlier results is caused by the inherent impreciseness of the automated annotation performed earlier. The earlier identified genes indeed encode enzymes related to lipid metabolism with some of the conserved domains present in enzymes required for fatty acid β-oxidation, but these conserved domains are also present in enzymes involved in other processes, such as lipid binding and biosynthetic thiolase reactions. The presence of these shared domains explains why these proteins were accidently annotated as being involved in β-oxidation of fatty acids (Berriman et al., 2009). Genes coding for acyl-CoA synthetases, on the other hand, are indeed present in the genome of *S. mansoni*, but these enzymes activate fatty acids by coupling them to coenzyme-A and the resulting acyl-CoAs are the beginning of several anabolic pathways that are known to be present in *S. mansoni* (Fig. 4).

As S. *mansoni* does not possess the enzymes necessary for fatty acid oxidation, how is it then possible that experiments seemed to indicate that fatty acid oxidation is essential for egg production by female *S. mansoni* (Huang et al., 2012). That study used inhibitors and RNAi to study fatty acid metabolism, while none of the experiments provides direct evidence for fatty acid oxidation, i.e. production of carbon dioxide from fatty acids. The first set of experiments showed that fecund female schistosomes use oxidative phosphorylation and that this process is essential for egg production. Oxidative phosphorylation by itself, however, is no indication for fatty acid oxidation. It has been shown earlier that in adult *S. mansoni* the degradation of glucose to carbon dioxide via Krebs cycle activity and oxidative phosphorylation produces at least one third of the ATP produced by glucose breakdown, the remainder is produced during the production of lactate, the major end product (Van Oordt et al., 1985). This explains why tampering with oxidative phosphorylation has a profound effect on energy consuming processes, such as egg laying. The second type of experiments was related to carnitine palmitoyl transferase 1 (CPT1, enzyme 2 in Fig. 4), an essential component of the pathway used for the import of fatty acids into mitochondria. This enzyme can be inhibited by etomoxir and it was observed that addition of this inhibitor resulted in a decrease of the oxygen consumption rate and in a decrease in the production of eggs in vitro (Huang et al., 2012). However, *S. mansoni* does not possess CPT1 (Fig. 4 and Fig. 5; Suppl Table 2) and etomoxir is known to inhibit not only CPT1 but also diacylglycerol acyltransferase (Xu et al., 2003), an anabolic enzyme catalyzing the final reaction in the synthesis of TAG, a process known to occur in schistosomes (Fig. 4). See below for a discussion on the importance of this enzyme for female schistosomes in the process of egg laying. In the third type of experiments RNAi and an inhibitor were used to study the effect of a decrease in activity of two enzymes on the rate of oxygen consumption and egg laying. The observation that inhibition or RNAi of acyl-CoA synthetase (enzyme 1 in Fig. 4) influences egg laying and oxygen consumption by female *S. mansoni* is not surprising in view of the importance of anabolic processes that occur during this energy consuming process. A correlation between activity of acyl-CoA activity and egg-laying is therefore no indication for fatty acid oxidation. The observed slight inhibition of egg laying after RNAi of acyl-CoA dehydrogenase (enzyme 4 in Fig 4) is enigmatic as schistosomes do not possess a gene for that enzyme (Fig. 4 and 5; Table 1; Suppl Table 2). As a fourth line of evidence for the occurrence and importance of fatty acid oxidation in egg-laying females, the presence of large lipid reserves in the vitellarium of fecund females is mentioned together with the observation that the decline of these lipid reserves correlates with egg laying and rate of oxygen consumption. These observations are, however, no indication for fatty acid oxidation and are the result of the use of lipids by female schistosomes to stuff the eggs. The role of lipids in the process of egg laying by the female and also during the further development of the egg is discussed below.

As fatty acids do not function in schistosomes as substrate for β-oxidation and therefore have no role as fuel in ATP production, the question raises what their real function in *S. mansoni* is. In general, fatty acids are present in cells mainly in two forms: they can be stored as triacylglycerols and they are used as building blocks of phospholipids and glycolipids. In this respect it should be realized that female worms produce approximately 350 eggs per day (Cheever et al., 1994) and each egg contains fatty acids present in triacylglycerol and phospholipids (Fig. 2). Hence, the female worm not only requires energy and amino acids for egg production, it also requires substantial amounts of fatty acids, and therefore, their daily uptake and digestion of red-blood cells is approximately 8 times higher than that of males (Cheever et al., 1994; Lawrence, 1973; Skelly et al., 2014). These ideas are in agreement with an earlier report of Newport and Weller (1982) who demonstrated that fatty acids are an absolute requirement for egg laying. The main lipid-constituent of the immature egg is triacylglycerol (Fig. 2), which is synthesized from diacylglycerol by the enzyme diacylglycerol acyltransferase (Smp_158510). Expression of this gene was shown to be strongly (10x) upregulated in fecund females in a bi-sex infection, in comparison to virgin females in a single sex infection (Anderson et al., 2015; Fitzpatrick et al., 2005; Lu et al., 2017). By whole mount *in situ* hybridization and RNA-seq it was shown that this enzyme is expressed exclusively in the vitellarium, which is the site of egg production (Wang and Collins, 2016). These results showed that female worms expel a large amount of fatty acids in lipids present in the excreted eggs and explains why fatty acids are essential for egg production. Fatty acids are essential components that, although they cannot be synthesized by the parasite itself, are used in anabolic processes. This explains why anabolic lipid metabolism in the female worm is crucial for egg production and why interference with it or with ATP generating processes such as oxidative phosphorylation will affect egg production (Huang et al., 2012).

As it was shown that fatty acids are essential for egg production and that interference with lipid metabolism affects egg production (Huang et al., 2012), we investigated the lipidome of eggs during maturation. As mature and immature eggs can be separated by density (Ashton et al., 2001), it is likely that a change in lipid composition occurs during egg development. Microscopic evaluation of developing eggs revealed that immature eggs contain many large lipid droplets (Fig. 1), which confirms earlier observations (Neill et al., 1988). Mature eggs, on the other hand, have far less lipid droplets and instead more cellular membranes were stained (Fig. 1). The lipidome of mature and immature eggs was then analyzed by LCMS, which demonstrated that mature eggs contain significantly more phospholipids than immature eggs. On the other hand, mature eggs contained less neutral lipids, albeit not significant. These observed changes in lipid composition of eggs prompted us to postulate that triacylglycerol in lipid droplets in the immature egg serves as a fatty acid storage that is used during egg maturation for phospholipid biosynthesis, which is needed for the formation of new membranes in the dividing cells of the developing miracidium.

In addition to the increase in phospholipid content, also the phospholipid composition changes during egg maturation. The lipidomic ‘finger-print’ of immature eggs was shown to be different from that of mature eggs, as indicated by the distinct and non-overlapping score plots in the PCA (Fig. 3A). The corresponding PCA loadings plot (Fig. 3B) showed that the difference between mature and immature eggs was reflected in many different lipid species from all phospholipid classes. These results showed that lipid metabolism is important during egg maturation as the lipid composition is substantially adjusted during development.

In conclusion, our results show that *S. mansoni* adults take up fatty acids and incorporate them in phospholipids and neutral lipids, such as triacylglycerol. Use of fatty acids by female schistosomes in these anabolic processes is crucial for egg production as the secreted eggs contain large amounts of fatty acids provided by the host. However, adult worms, eggs and miracidia do not oxidize fatty acids, and therefore, fatty acid catabolism does not occur and thus does not contribute to ATP production. Genome analysis showed that previously postulated genes for fatty acid β-oxidation were incorrectly annotated or attributed and that *S. mansoni* in fact lacks genes encoding the enzymes required for fatty acid β-oxidation. Hence, *S. mansoni* does not and cannot oxidize fatty acids (nor can *S. japonicum*).

## Footnotes

1 After writing this manuscript we happened to notice the announcement of a new analysis of all available genomes of helminths. In that analysis it is also concluded that schistosomes indeed lack the enzymes necessary for β-oxidation of fatty acids (Coglan et al.; cited by Jex et al. Trends in Parasitology, in press, available on line 24 October 2018). However, in that genomic study by Coglan et al. it is mentioned that: “However, biochemical data suggest they do perform β-oxidation, so they may have highly diverged but functional β-oxidation”. Our experiments, however, provided no indication whatsoever that schistosomes can perform β-oxidation of fatty acids. Furthermore, our in-depth genomic analysis, specifically of the highly conserved domains in the various β-oxidation enzymes, showed that *S. mansoni* lacks genes for the enzymes of β-oxidation of fatty acids, but possesses true acyl-CoA synthetases, and clarifies why earlier annotations were incorrect.

